# Myosin ATPase inhibition fails to rescue the metabolically dysregulated proteome of nebulin-deficient muscle

**DOI:** 10.1101/2024.05.07.592906

**Authors:** Jenni Laitila, Robert A.E. Seaborne, Natasha Ranu, Justin S. Kolb, Carina Wallgren-Pettersson, Nanna Witting, John Vissing, Juan Jesus Vilchez, Edmar Zanoteli, Johanna Palmio, Sanna Huovinen, Henk Granzier, Julien Ochala

## Abstract

Nemaline myopathy (NM) is a genetic muscle disease, primarily caused by mutations in the *NEB* gene (*NEB-*NM) and with muscle myosin dysfunction as a major molecular pathogenic mechanism. Recently, we have observed that the myosin biochemical super-relaxed state was significantly impaired in *NEB*-NM, inducing an aberrant increase in ATP consumption and remodelling of the energy proteome in diseased muscle fibres. As the small-molecule Mavacamten is known to promote the myosin super-relaxed state and reduce the ATP demand, here, we tested its potency in the context of *NEB*-NM. We first conducted *in vitro* experiments in isolated single myofibres from patients and found that Mavacamten successfully reversed the myosin ATP over-consumption. Following this, we assessed its short-term *in vivo* effects by using the conditional nebulin knock-out (c*Neb* KO) mouse model and by subsequently performing global proteomics profiling in dissected soleus myofibres. After a four-week treatment period, we observed a remodelling of a large number of proteins in both c*Neb* KO mice and their wild-type siblings. Nevertheless, these changes were not related to the energy proteome, indicating that short-term Mavacamten treatment is not sufficient to properly counterbalance the metabolically dysregulated proteome of c*Neb* KO mice. Taken together, our findings emphasize Mavacamten potency *in vitro* but challenge its short-term efficacy *in vivo*.

**Key points summary:**

- No cure exists for nemaline myopathy, a type of genetic skeletal muscle disease mainly derived from mutations in genes encoding myofilament proteins.
- Applying Mavacamten, a small molecule directly targeting the myofilament, to isolated membrane-permeabilized muscle fibres from human patients restored myosin energetic disturbances.
- Treating a mouse model of nemaline myopathy *in vivo* with Mavacamten for four weeks, remodeled the skeletal muscle fibre proteome without any noticeable effects on energetic proteins.
- Short-term Mavacamten treatment may not be sufficient to reverse the muscle phenotype in nemaline myopathy.

## Introduction

Nemaline myopathy (NM) is a congenital myopathy, with an estimated incidence of 1 in 50,000 live births (Jungbluth *et al*., 2018; Laitila & Wallgren-Pettersson, 2021). Its clinical symptoms are heterogenous in terms of presentation, severity and progression. Nevertheless, the typical muscle phenotype consists of hypotonia, weakness, fatigue and leanness (Jungbluth *et al*., 2018; Laitila & Wallgren-Pettersson, 2021). At least twelve causative genes have been identified to date, with *NEB* mutations accounting for more than 50% of all NM cases - *NEB*-NM (Jungbluth *et al*., 2018; Fisher *et al*., 2022). These *NEB* mutations often result in partial or (in rare cases) complete deficiency of a protein named nebulin in skeletal muscle (Jungbluth *et al*., 2018; Laitila & Wallgren-Pettersson, 2021). This induces a cascade of molecular and cellular pathophysiological events by which myosin motors cannot bind properly to actin filaments, preventing optimal force-producing capacity at the level of the myofibre, overall significantly contributing to hypotonia and muscle weakness in *NEB*-NM (Ottenheijm *et al*., 2009; Ochala *et al*., 2011; Lindqvist *et al*., 2016; Ross *et al*., 2019). Based on this, myosin has been targeted in human cells/fibres and in animal preclinical models of *NEB*-NM. We, and others, have then demonstrated that hyper-activating this specific motor protein, pharmacologically or through recombinant adeno-associated virus-related gene therapy, partially alleviates muscle weakness (Lindqvist *et al*., 2016; Lindqvist *et al*., 2019; Ross *et al*., 2019). However, these studies have failed to fully rescue the *in vivo* muscle phenotype, emphasizing the complexity of *NEB*-NM and the limited efficacy of myosin activators.

Myosin molecules have two known relaxed biochemical states, i.e., ‘super-relaxed’ (SRX) and ‘disordered-relaxed’ (DRX) states (Stewart *et al*., 2010; McNamara *et al*., 2015). SRX and DRX states mainly differ in their ATP usage, with the DRX state having an ATPase activity five to ten times higher than the SRX state (Stewart *et al*., 2010; McNamara *et al*., 2015). Following this, we have recently shown that, in human fibres and in a mouse model of *NEB*-NM, the proportion of myosin molecules in the energy-demanding DRX state is surprisingly greater than normal, leading to an unusually high basal skeletal muscle ATP consumption (Ranu *et al*., 2022). We have also shown that such accrued demand is accompanied by remodelling of the energy proteome, likely contributing to muscle fatigue and leanness in *NEB*-NM (Ranu *et al*., 2022). In theory, applying an intervention that hyper-activates myosin pharmacologically or genetically as performed in previous studies (Lindqvist *et al*., 2016; Lindqvist *et al*., 2019; Ross *et al*., 2019), would worsen such a pathogenetic metabolic mechanism. Hence, in the present study, we aimed to test the potency of therapeutically inhibiting myosin in the context of *NEB*-NM.

Among myosin inhibitors is Mavacamten (also known as MYK-461 or Camzyos), a selective allosteric modulator that inhibits the ATP turnover of myosin heads by stabilizing the SRX state, functionally reducing the number of myosin molecules in the DRX state and, ultimately, lowering the basal resting ATP demand of striated muscle (Anderson *et al*., 2018; Gollapudi *et al*., 2021). When Mavacamten is administered to patients with hypertrophic cardiomyopathy, the proportion of myosin heads in the DRX state is reduced, relieving cardiac symptoms, and increasing quality of life and functional status (Olivotto *et al*., 2020). Thus, here, we specifically aimed to test the efficacy of Mavacamten for *NEB*-NM in skeletal muscle. We initially hypothesized that Mavacamten would counterbalance the disproportionate high level of myosin molecules in the DRX state, allowing a rescue of the muscle energy proteome.

## Materials and Methods

### Human subjects

Included in the study were upper leg muscle biopsy specimens from twelve NM patients with *NEB* mutations (*NEB*-NM) and 12 age-and gender-matched controls with no history of neuromuscular disease. All tissue was consented, stored, and used in accordance with the Human Tissue Act, UK, under local ethical approval (REC 13/NE/0373). Details of all the subjects are presented in Table 1. All samples were snap-frozen and stored at -80°C until used.

**Table 1:** Patient and control muscle biopsy samples used.

| Age (years) | Gender | Mutation | Disease | Source |
| --- | --- | --- | --- | --- |
| 70 | Female | *NEB* (c.22594C>T and c.5238+335_5764-407del) | Nemaline myopathy | Helsinki, Finland |
| 28 | Female | *NEB* (c.2836-2A>G and c.5763+5G>A) | Nemaline myopathy | Copenhagen, Denmark |
| 46 | Male | *NEB* (c.17234C>T and c.2271_22713del) | Nemaline myopathy | Copenhagen, Denmark |
| 32 | Male | *NEB* (c.23526_23527del and c.24792_24793del) | Nemaline myopathy | Helsinki, Finland |
| 22 | Male | *NEB* (c.12667G>A) | Nemaline myopathy | Valencia, Spain |
| 70 | Female | *NEB* (c.19097G>T and c.508-7T>A) | Nemaline myopathy | Helsinki, Finland |
| 22 | Male | *NEB* (c.18164T>C and c.15973C>T) | Nemaline myopathy | Valencia, Spain |
| 9 | Male | *NEB* (c.12676_12677delTA and c.10357A>C) | Nemaline myopathy | Valencia, Spain |
| 10 | Female | *NEB* (c.4489A>G and c.11947_11948 in sAGGACTATG) | Nemaline myopathy | Sao Paulo, Brazil |
| 7 | Male | *NEB* (c.19101+5G>A and c.9723+1G>A) | Nemaline myopathy | Sao Paulo, Brazil |
| 9 | Female | *NEB* (c.19102-4del and c.8889+1G>A) | Nemaline myopathy | Sao Paulo, Brazil |
| 25 | Female | *NEB* (Deletion of exons 11 to 107) | Nemaline myopathy | Tampere, Finland |
| 55 | Male | - | - | London, UK |
| 25 | Female | - | - | London, UK |
| 63 | Female | - | - | London, UK |
| 37 | Male | - | - | London, UK |
| 44 | Female | - | - | London, UK |
| 38 | Female | - | - | Copenhagen, Denmark |
| 45 | Female | - | - | Copenhagen, Denmark |
| 49 | Male | - | - | Copenhagen, Denmark |
| 21 | Female | - | - | Copenhagen, Denmark |
| 24 | Female | - | - | Copenhagen, Denmark |
| 35 | Female | - | - | Copenhagen, Denmark |
| 41 | Female | - | - | Copenhagen, Denmark |

### Nebulin knockout mouse model and Mavacamten treatment

The conditional muscle-specific nebulin knockout mouse model used in the study has previously been published in detail (Li *et al*., 2015). Briefly, mice were on a C57BL/6J background. Floxed mice were bred to a MCK-Cre strain that expresses Cre recombinase under the control of the Muscle Creatine Kinase (MCK) promoter. Mice that were positive for MCK-Cre and homozygous for the floxed nebulin allele were nebulin deficient (cNeb KO). Mice with one nebulin wild-type allele and being either MCK-Cre positive or negative served as controls. All experiments were approved by the University of Arizona Institutional Animal Care and Use Committee (09-056) and were in accordance with the United States Public Health Service’s Policy on Humane Care and Use of Laboratory Animals. At week 8, cNeb KO and wild-type female mice were subjected to a short-term Mavacamten treatment. Mavacamten was added to the drinking water (hydropack) and administered at a safe and efficient dose of 2.5 mg.kg^-1^.day^-1^ for four weeks (Toepfer *et al*., 2019). At week 12, animals were weighed, anesthetized with isoflurane, and sacrificed by cervical dislocation. Soleus skeletal muscles were then dissected and flash-frozen in liquid nitrogen before being stored at -80°C for later analysis.

### Solutions

As previously published (Ross *et al*., 2019), the relaxing solution contained 4 mM Mg-ATP, 1 mM free Mg^2+^, 10^-6.00^ mM free Ca^2+^, 20 mM imidazole, 7 mM EGTA, 14.5 mM creatine phosphate and KCl to adjust the ionic strength to 180 mM and pH to 7.0. Additionally, the rigor buffer for Mant-ATP chase experiments contained 120 mM K acetate, 5 mM Mg acetate, 2.5 mM K_2_HPO_4_, 50 mM MOPS, 2 mM DTT with a pH of 6.8.

### Muscle preparation and fibre permeabilisation

Cryopreserved human and mouse muscle samples were immersed in a membrane-permeabilising solution (relaxing solution containing glycerol; 50:50 v/v) for 24 hours at -20°C, after which they were transferred to 4°C and bundles of approximately 50 to 100 muscle fibres were dissected free.

These bundles were kept in the membrane-permeabilising solution at 4°C for an additional 24 hours (to allow for a proper skinning/membrane permeabilisation process). After these steps, bundles were stored in the same buffer at -20°C for use up to one week (Ross *et al*., 2019; Ross *et al*., 2020).

### Mant-ATP chase experiments

On the day of the experiments, bundles were transferred to relaxing solution and single myofibres were manually isolated. Their ends were individually clamped to half-split copper meshes designed for electron microscopy (SPI G100 2010C-XA, width, 3 mm), which had been glued to glass slides (Academy, 26 x 76 mm, thickness 1.00-1.20 mm). Cover slips were then attached to the top (using double-sided tape) to create flow chambers (Menzel-Glaser, 22 x 22 mm, thickness 0.13-0.16 mm) (Ochala *et al*., 2021). Muscle fibres were mounted at a slack length (with their sarcomere length checked using the brightfield mode of a Zeiss Axio Scope A1 microscope, approximately at 2.20 µm). Similarly to previous studies (Ochala *et al*., 2021), all experiments were performed at 25°C, and each fibre was first incubated for five minutes with a rigor buffer. A solution containing the rigor buffer with 250 μM Mant-ATP was then flushed and kept in the chamber for five minutes. At the end of this step, another solution made of the rigor buffer with 4 mM unlabelled ATP was added with simultaneous acquisition of the Mant-ATP chase.

For fluorescence acquisition, a Zeiss Axio Scope A1 microscope was used with a Plan-Apochromat 20x/0.8 objective and a Zeiss AxioCam ICm 1 camera. Frames were acquired every five seconds for the first 90 seconds and every 10 seconds for the remaining time with a 20 ms acquisition/exposure time using a DAPI filter set, and images were collected for five minutes. Three regions of each individual myofibre were sampled for fluorescence decay using the ROI manager in ImageJ as previously published (Ochala et al., 2021). The mean background fluorescence intensity was subtracted from the average of the fibre fluorescence intensity (for each image taken). Each time point was then normalized by the fluorescence intensity of the final Mant-ATP image before washout (T = 0). These data were then fit to an unconstrained double exponential decay using Graphpad Prism 9.0:

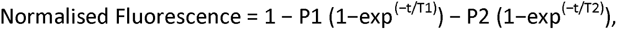

where P1 is the amplitude of the initial rapid decay approximating the disordered-relaxed state with T1 as the time constant for this decay. P2 is the slower second decay approximating the proportion of myosin heads in the super-relaxed state with its associated time constant T2 (Ochala *et al*., 2021).

### Single muscle fibre force-generating capacity

As for Mant-ATP experiments, individual myofibres were dissected in the relaxing solution. They were then individually attached between connectors leading to a force transducer (model 400A; Aurora Scientific) and a lever arm system (model 308B; Aurora Scientific). Sarcomere length was set to ≈2.50 µm and the temperature to 15°C (Ochala *et al*., 2021). As the baths had glass bottom and right-angle prisms, fibre cross-sectional area could be estimated from the width and depth, assuming an elliptical circumference. To determine the maximum isometric force generating capacity, myofibres were bathed in an activating buffer (pCa 4.5). Specific force corresponded to absolute force normalized to myofibre cross-sectional area.

### Immunofluorescence staining and imaging

To avoid any potential misinterpretation due to the type of myosin heavy chain, for the human/mouse Mant-ATP chase experiments or cellular force measurements, we assessed the sub-type using immunofluorescence staining as previously described (Ochala *et al*., 2021). Briefly, flow-chamber mounted myofibres were stained with an anti-β-cardiac/skeletal slow myosin heavy chain antibody (IgG1, A4.951, sc-53090 from Santa Cruz Biotechnology, dilution: 1:50). Myofibres were then washed in PBS/0.025% Tween-20 and incubated with secondary antibodies: goat anti-mouse IgG1 Alexa 555 (from ThermoScientific, dilution 1:1,000), in a blocking buffer. After washing, muscle fibres were mounted in Fluoromount. To identify the type of fibres, images were acquired using a confocal microscope (Zeiss Axiovert 200, 63x oil objective) equipped with a CARV II confocal imager (BD Biosciences) (Ross *et al*., 2019; Ross *et al*., 2020).

### Mouse sectioning and CSA measurements

Immunolabelling was performed on 10 μm cryosections, fixed in 4% PFA (10 min), permeabilized in 0.1% Triton X-100 (20 min) and blocked in 10% Normal Goat Serum (50062Z, Life Technologies) with 0.1% BSA (1 h). Sections were incubated o/n (4°C) with primary antibody against Laminin (Rabbit polyclonal, Sigma, L9393, diluted 1:20). Alexa Fluor Donkey anti-Rabbit 488 (A11034) was used as the secondary (Life Technologies, 1:500 in 10% Normal Goat Serum). For fibre typing, Zenon™ Mouse IgG1 Labelling Kits with Alexa Fluor 488 (Z25002) and Alexa Fluor 594 (Z25007) were used for labelling MYH I (mouse monoclonal A4.951, DSHB, 1:25) and MYH IIA (mouse monoclonal SC71, DSHB, 1:25). The labelling complexes were added in 5% goat serum with 0.1% BSA and 0.1% Triton X-100 and incubated for 2h RT. Fluorescent images were obtained with a 10x objective on a Zeiss Axio Observer 3 fluorescence microscope with a Colibri 5 led detector, combined with Zeiss Axiocam 705 mono camera, using Zen software (Zeiss) and exported to single channel .tiff files. The single images with fibre membrane staining were then loaded into ilastik (version 1.4.0.post1, www.ilastik.org) and the software was trained to distinguish between muscle fibres and their membrane. The results were exported from ilastik as binary H5 files with the Simple Segmentation setting and analyzed in Fiji (ImageJ version 1.53c with ilastik Fiji plugin, https://imagej.net/) with a macro using the analysis particle function and ROI manager tool (region of interest). Briefly, the macro automatically generated a ROI-set with individual ROI’s outlining the perimeter of the muscle fibres. The ROI-set was then visually inspected and errors were manually corrected, before the cross-sectional areas were exported to an Excel file. Fibre type was manually assigned to the unique ID of each ROI in the Excel file (Schindelin *et al*., 2012; Berg *et al*., 2019).

### Global proteome profiling of mouse skeletal muscle

As the skeletal muscle niche contains an array of cell types that may lead to non-myofibre specific data sets when analysing bulk tissue (Seaborne & Ochala, 2023), we enriched for myofibers in our mouse proteomic experiments. Dissected soleus muscle specimens (N=4 per condition) were retrieved from the -80°C freezer and placed in 0.2% collagenase at 37°C for 60-90 minutes and routinely disturbed to enable disruption. Samples were transferred to a sterilised 6-well plate, agitated to further dissociate the sample and individual myofibres were manually/pipette transferred to ice-cold PBS under a dissection microscope. An estimated 50 myofibres were withdrawn from 6-well plate, washed through with PBS for a second time, snap-frozen on dry-ice and stored at -80°C. These were then processed as in (Lewis *et al*., 2023). Briefly, they were lysed with lysis buffer (1% (w/v) Sodium Deoxycholate, 100 mM Teab, pH 8.5) and incubated for 10 min at 95°C followed by sonication using a Bioruptor pico (30 cycles, 30 sec on/off, ultra-low frequency). Heat incubation and sonication were repeated once, samples cleared by centrifugation, reduced with 5 mM (final concentration) of TCEP for 15 min at 55°C, alkylated with 20 mM (final concentration) CAA for 30 minutes at RT, and digested adding Trypsin/LysC at 1:100 enzyme/protein ratio. Peptides were cleaned up using StageTips packed with SDB-RPS and resuspended in 50 µL TEAB 100 mM, pH 8,5. 50 µg of each sample was labelled with 0.5 mg TMTpro labelling reagent according to the manufacturer’s instructions. Labelled peptides were combined and cleaned up using C18-E (55 µm, 70 Å, 100 mg) cartridges (Phenomenex). Labelled desalted peptides were resuspended in buffer A* (5% acetonitrile, 1% TFA), and fractionated into 16 fractions by high-pH fractionation. For this, 20 ug peptides were loaded onto a Kinetex 2.6u EVO C18 100 Å 150 x 0.3 mm column via an EASY-nLC 1200 HPLC (Thermo Fisher Scientific) in buffer AF (10 mM TEAB), and separated with a non-linear gradient of 5 – 44 % buffer BF (10mM TEAB, 80 % acetonitrile) at a flow rate of 1.5 µL / min over 62 min.

Fractions were collected every 60 s with a concatenation scheme to reach 16 final fractions (e.g. fraction 17 was collected together with fraction 1, fraction 18 together with fraction 2, and so on). Fractions were evaporated, resuspended in buffer A*, and measured on a Vanquish Neo HPLC system (Thermo Fisher Scientific) coupled through a nano-electrospray source to a Tribrid Ascend mass spectrometer (Thermo Fisher Scientific). Peptides were loaded in buffer A (0.1 % formic acid) onto a 110 cm mPAC HPLC column (Thermo Fisher Scientific) and separated with a non-linear gradient of 1 – 50 % buffer B (0.1 % formic acid, 80 % acetonitrile) at a flow rate of 300 nL/min over 100 min. The column temperature was kept at 50° C. Samples were acquired using a Real Time Search (RTS) MS3 data acquisition where the Tribrid mass spectrometer was switching between a full scan (120 K resolution, 50 ms max. injection time, AGC target 100%) in the Orbitrap analyzer, to a data-dependent MS/MS scan in the Ion Trap analyzer (Turbo scan rate, 23 ms max. injection time, AGC target 100% and HCD activation type). Isolation window was set to 0.5 (m/z), and normalized collision energy to 32. Precursors were filtered by charge state of 2-5 and multiple sequencing of peptides was minimized by excluding the selected peptide candidates for 60 s. MS/MS spectra were searched in real time on the instrument control computer using the Comet search engine with either the UP000291022 U. americanus or UP000005215 *I. tridecemlineatus* FASTA file, 0 max miss cleavage, 1 max oxidation on methionine as variable mod. and 35 ms max search time with an Xcorr soring threshold of 1.4 and 20 precursor ppm error). MS/MS spectra resulting in a positive RTS identification were further analyzed in MS3 mode using the Orbitrap analyzer (45K resolution, 105 ms max. injection time, AGC target 500%, HCD collision energy 55 and SPS = 10). The total fixed cycle time, switching between all 3 MS scan types, was set to 3 s.

### Proteome profiling analysis

Analyses of all protein expression data was performed with the automated analysis pipeline of the Clinical Knowledge Graph, as previously detailed (Santos *et al*., 2022). Briefly, LFQ intensities were Log2 transformed, with proteins with less than at least 2 valid values in at least one comparative group filtered out. Missing values were imputed using a mixed imputation method. That is, on a per protein bases, missing values in samples belonging to the same comparative group were KNN imputed if at least 60% valid values existed in that same group. Thereafter, remaining missing values were imputed using the MinProb approach (width = 0.3, shift = 1.8), as previously described (Lazar *et al*., 2016). Unpaired T-test comparisons across relevant conditions were performed in Perseus (Tyanova & Cox, 2018), with differentially expressed proteins considered statistically significant following 250-permutation based FDR correction, at a level of 0.05 and FC = 1, unless otherwise stated. Significant proteins were used for functional enrichment analyses alongside relevant Gene

Ontology (GO) annotations, with a Fisher’s exact test and Benjamini-Hocheberg correction applied (> 2 hits per term), as per the Clinical Knowledge Graph pipeline. Significantly enriched GO terms for analysis of the top 250 proteins from all data sets (e.g. Fig 4C), ShinyGO (v.0.77 (Ge *et al*., 2020)) were used with background set as all identified proteins within the data set. Significant GO terms were extracted, GO IDs retrieved using the pRoloc in R (v. 12.0), and packages REVIGO (Supek *et al*., 2011), MetaScape (Zhou *et al*., 2019) and cystoscope (v. 3.9.1) were used, alongside adjusted significance values, to generate association networks. Log2 transformed LFQ values for all myosin isoforms were extracted and an ANOVA with Tukey post-hoc was used to compare across conditions (R studio, v12.0).

### Statistical analysis

Data are then presented as means ± standard deviations. For all non-protein data sets, graphs were prepared and analysed in Graphpad Prism v9. Statistical significance was set to P < 0.05. T-tests or various ANOVAs with Tukey post-hoc were run to compare groups/treatment (Ross *et al*., 2019). Statistical analyses for protein data sets are detailed above.

## Results

### Mavacamten restores myosin relaxed states in humans and mice

We started by evaluating the effects of Mavacamten on the fraction of myosin heads in the ATP-demanding DRX and ATP-conserving SRX conformations in membrane-permeabilised muscle fibres derived from twelve human controls and twelve *NEB*-NM patients (Table 1). As *NEB*-NM patients have a high predominance of muscle fibres expressing the β/slow myosin heavy chain (Jungbluth *et al*., 2018), we only applied the Mant-ATP chase assay on this particular fibre type (Ranu *et al*., 2022). A total of 181 individual muscle fibres were exposed to four different concentrations of Mavacamten: 0, 1, 5 and 10 µM (6 to 8 myofibres for each of the 24 subjects - Table 1). Interestingly, even a small concentration of Mavacamten (1 µM) was sufficient to significantly decrease the amount of myosin molecules in the DRX state in *NEB*-NM patients (P1 and P2 - Fig. 1A-C). Such decrease was further pronounced at greater concentrations of Mavacamten in both controls and *NEB*-NM patients (5 and 10 µM - Fig. 1A-C). However, the actual ATP turnover time of myosin heads in the DRX and SRX conformations was not altered, even though a dose-dependent gradual trend was noticed (T1 and T2 - Fig. 1D-E). To further assess whether these changes related to Mavacamten would impact muscle fibre function as previously suggested (Scellini *et al*., 2021), we measured the cellular maximum force-generating capacity at the same four concentrations of Mavacamten: 0, 1, 5 and 10 µM. A total of 142 myofibres expressing the β/slow myosin heavy chain was evaluated (5 to 7 myofibres for each of the 24 subjects - Table 1). Even though *NEB*-NM patients had significantly lower forces than controls, we did not detect any effect of Mavacamten on force at any concentration (Fig. 1F).

**Figure 1:**
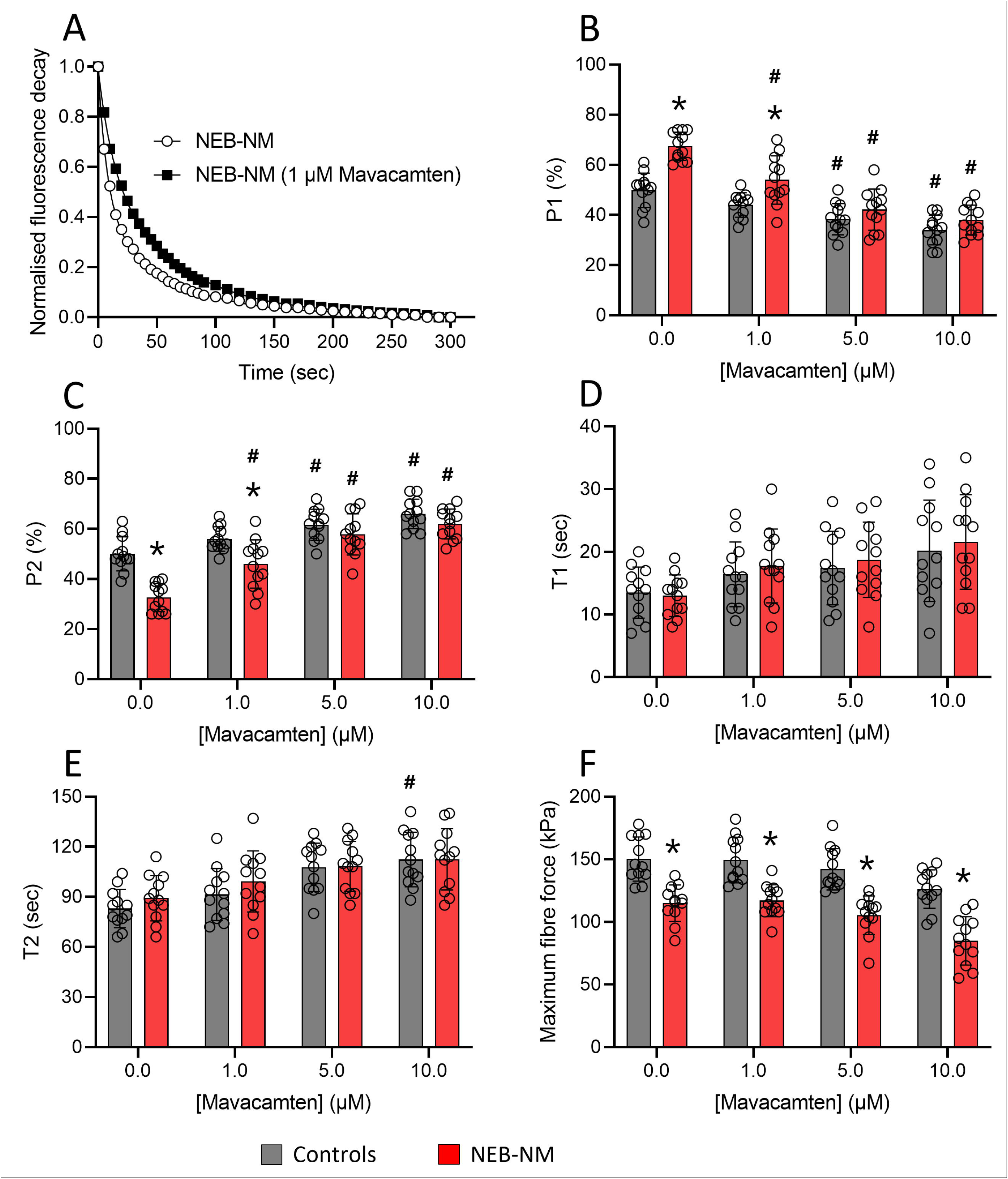
Mavacamten alters the proportions of myosin heads in the DRX and SRX states in controls and. NEB**-NM patients.** Typical Mant-ATP chase experimental data show exponential decays for one individual myofibre without and with Mavacamten (1 µM). The proportion of myosin molecules in the DRX: disordered-relaxed (P1, B) and SRX: super-relaxed states (P2, C) as well as their respective ATP turnover lifetimes (T1, D and T2, E) with varying concentrations of Mavacamten are displayed. (F) shows the cellular force generating capacity as a function of Mavacamten concentration. Dots are individual subject’s average data. Means and standard deviations also appear on histograms. * denotes a difference (P < 0.05) when compared with controls. ^#^ denotes a difference when compared with no Mavacamten for the same group of subjects (either controls or *NEB*-NM patients).

We decided to validate the above human results in an animal model of *NEB*-NM, before testing Mavacamten *in vivo*. Based on the fact that most of the *NEB*-NM patients survive to adulthood with symptoms including hypotonia, weakness and fatigue, we focused on a conditional nebulin KO mouse model (c*Neb* KO) where muscle-specific deletions are present from birth and where symptoms mimic the human NM condition (Li *et al*., 2015). We first performed the Mant-ATP chase experiments and separated the results depending on the myosin heavy chain expressed by the muscle cells, either β/slow or type II. 45 soleus myofibres expressing the β/slow myosin heavy chain (5 to 6 myofibres from each of the four c*Neb* KO mice and four wild-type siblings) and 70 type II muscle fibres (7 to 10 myofibres per animal) were subjected to increasing concentrations of Mavacamten: 0, 1, 5 and 10 µM. In muscle fibres expressing the β/slow myosin heavy chain, as in humans, at 1 µM of Mavacamten, there was a significant decrease in the proportion of myosin heads in the DRX state and increase in the SRX state in c*Neb* KO mice (P1 and P2 - Fig. 2A-B). This effect was exacerbated in the presence of 5 and 10 µM of Mavacamten in both c*Neb* KO mice and wild-type fibres (Fig. 2A-B). The ATP turnover time of myosin molecules in the DRX and SRX conformations (T1 and T2 - Fig. 2C-D) as well as the maximum isometric force production (Fig. 2E) were, however, not affected by Mavacamten at any concentration. In type II myofibres, we did not observe a significant difference in any of the parameters (P1, P2, T1, T2 and force). We only noticed a tendency towards a lower fraction of myosin molecules in the DRX state at 5 and 10 µM of Mavacamten (Fig. 2A-F).

**Figure 2:**
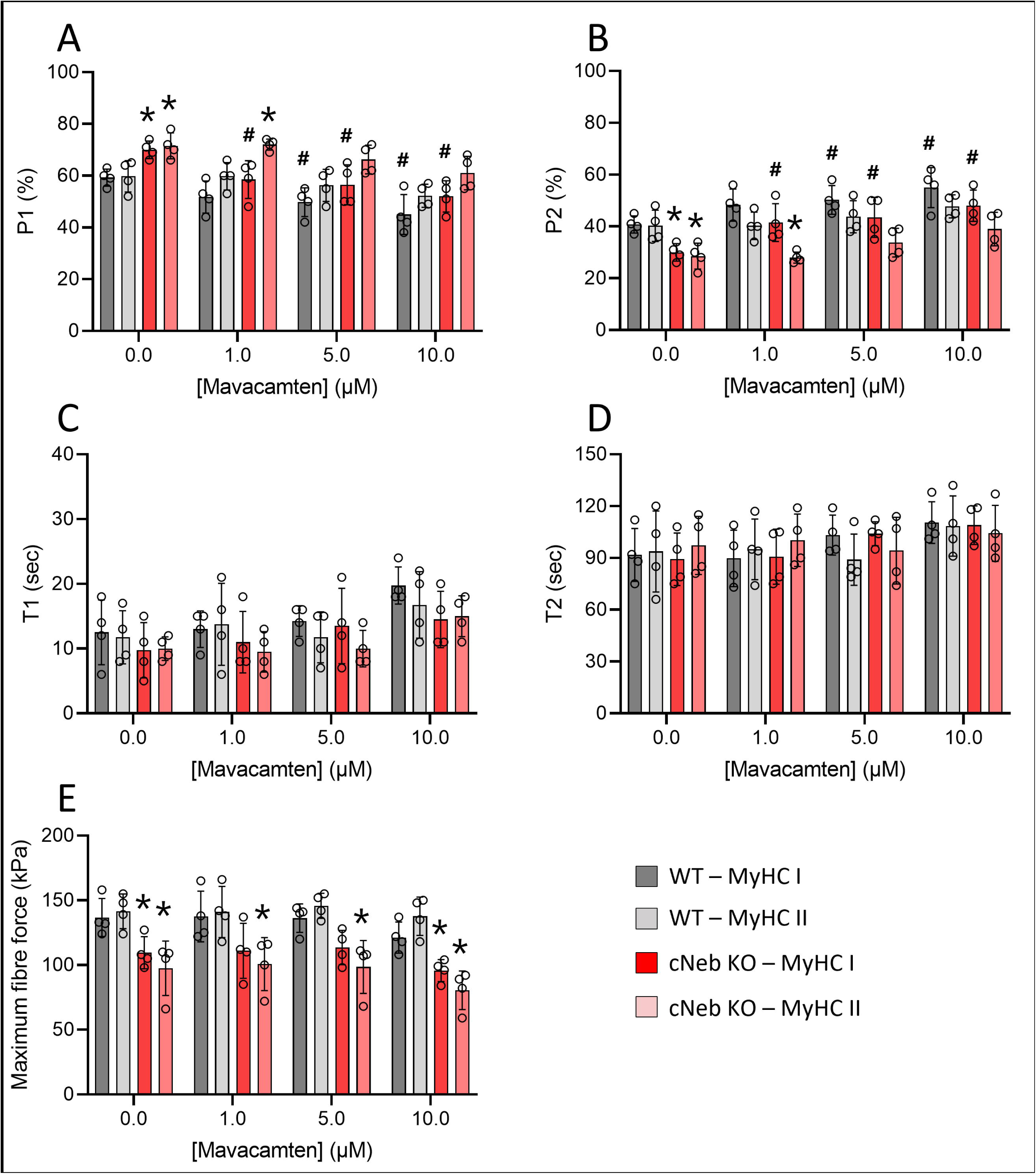
Mavacamten modifies the fraction of myosin molecule in the DRX and SRX states in cNeb KO mice. The fraction of myosin heads in the DRX: disordered-relaxed (P1, A) and SRX: super-relaxed states (P2, B) as well as their respective ATP turnover lifetimes (T1, C and T2, D) with increasing concentrations of Mavacamten are presented. (E) displays the myofibre maximum isometric force production at various Mavacamten concentrations. Dots are individual mouse’s average data. Means and standard deviations also appear on histograms. * denotes a difference (P < 0.05) when compared with wild-type siblings. ^#^ denotes a difference when compared with no Mavacamten for the same group of animals (either wild-type or transgenic).

Altogether, our data indicate that, in resting skeletal myofibres of controls as well as in *NEB*-NM patients/c*Neb* KO mice, small doses of Mavacamten are potent, lowering the basal ATP consumption without causing any major negative impact on the contractile maximal force.

### Mavacamten treatment remodels the proteome in mice

In line with our findings on c*Neb* KO mice, we sought to determine whether Mavacamten would be a potent small molecule *in vivo* by lowering the ATP consumption and acting on muscle metabolism (Jungbluth *et al*., 2018; Laitila & Wallgren-Pettersson, 2021). For that, 8-week-old c*Neb* KO mice and wild-type siblings were exposed to vehicle (DMSO) or Mavacamten for a short period of time (For 4 weeks, 2.5 mg.kg^-1^.day^-1^ administered in drinking water, known to be safe and efficient as in another published work (Toepfer *et al*., 2019)). After four weeks, mice treated with Mavacamten had no significant differences in their body weight compared with rodents treated with the vehicle (Fig. 3A). Similarly, the normalised body weight gains were not significantly different even though a tendency towards a greater weight gain was noticed for c*Neb* KO mice treated with Mavacamten compared with animals treated with the vehicle (Fig. 3B). To gain insights into any potential subtle muscle-specific changes, we dissected the soleus muscles of all the animals from the four groups. We specifically chose this skeletal muscle as it is one of the few containing a mixture of muscle fibres with either β/slow or type II myosin heavy chains as in humans. We did not find any significant difference in their absolute or normalised (to body weight) muscle mass (Fig. 3C-D) nor normalised tibia length to body weight (Fig. 3E). Nevertheless, and unexpectedly, fibre cross-sectional area was significantly smaller in type I and II fibres of c*Neb* KO mice treated with Mavacamten compared with fibres of transgenic rodents treated with the vehicle (Fig. 3F). Finally, a treatment-related trend towards a faster profile was noticed when analysing myofibre type proportions (Fig. 3G-H).

**Figure 3:**
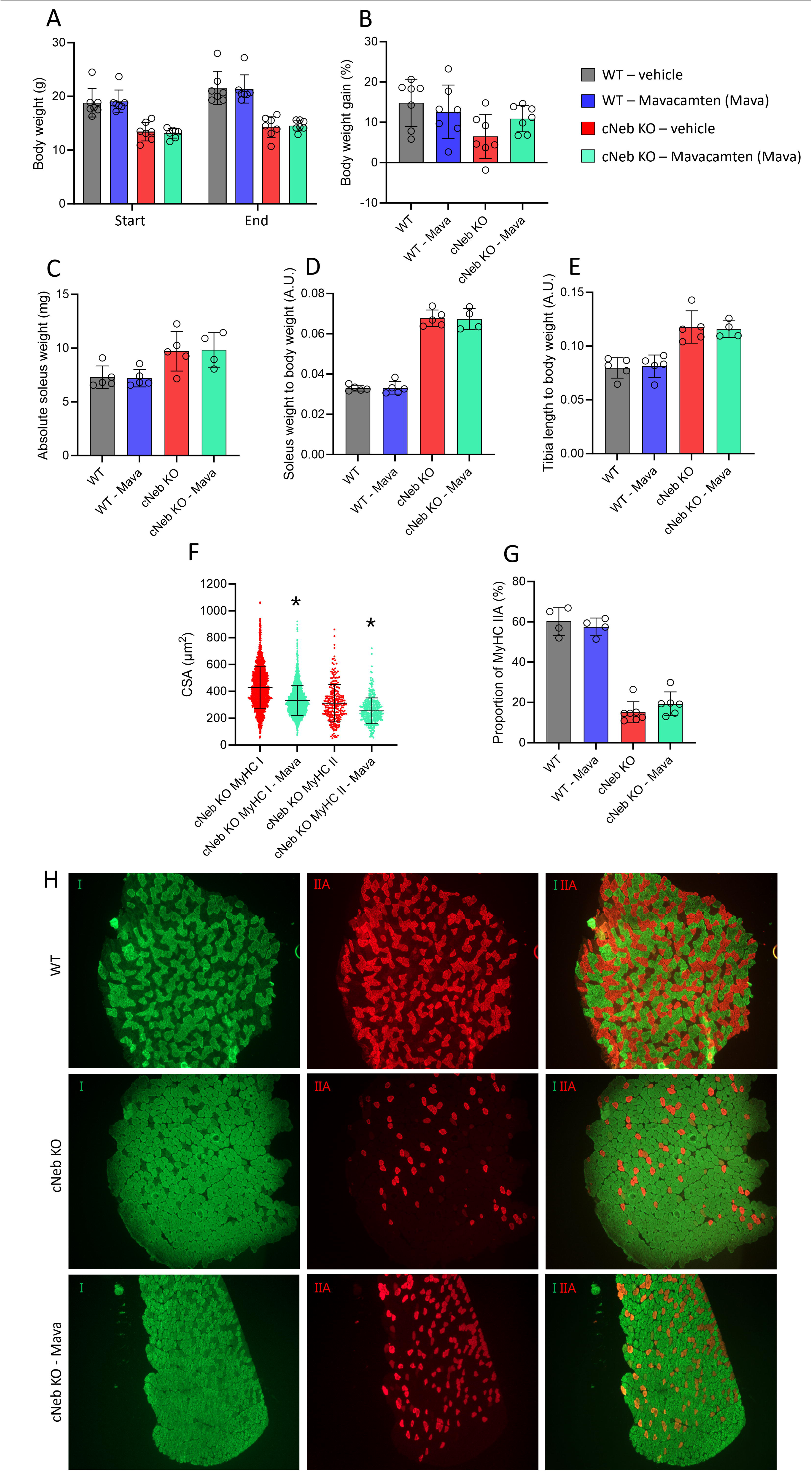
Effects of a Mavacamten treatment on body weight and muscle fibre type proportions in cNeb KO mice. Body weight (A), body weight gain (B), absolute soleus muscle wet weight (C), soleus wet weight to body weight (D), tibia length to body weight (E), fibre cross-sectional area (F) and proportion of muscle fibre expressing the type IIA myosin heavy chain isoform (G) are presented. Typical images allowing us to calculate the myosin heavy chain composition are also displayed (H). Dots are individual mouse’s data. Means and standard deviations also appear on histograms. * denotes a difference (P < 0.05) when compared with a vehicle treatment for the same group of animals (either wild-type or transgenic).

To explore potential differences at the protein level, we enriched myofibres from the soleus muscles and performed global proteomic profiling. These data sets yielded a large number of targets across all conditions (Fig. 4A) with high specificity towards proteins either heavily enriched or specific to myofibres (Fig. 4B-C). Our initial comparison focussed on c*Neb* KO vs wild-type siblings (WT) to establish the extent of dysregulation associated with the disease model. Here, we found a significant difference in the global proteomic profile of these conditions, as highlighted by clustering on principle component analyses (Fig. 4D), where disease state makes up the majority of the differences. This translated into > 1500 protein targets showing differential expression, with an almost 4-fold increase in the number of proteins up-regulated in c*Neb* KO compared with those down-regulated (Fig. 4E; Supp. File 1 & 2). Unsurprisingly, we observed an increase in proteins associated with ubiquitination (Fig. 4F-G), including FBXO40, TRIM35, HDAC3 and FBXO25. Interestingly. however, it was the proteins that were downregulated that seemed to provide the most coordinated biological response. Indeed, 9 of the top 10 most significantly enriched GO biological process terms (hits < 2, FRD < 0.05; Supp. File 3) were directionally down-regulated in the c*Neb* KO model (Fig. 4E). These GO terms cluster into small networks largely revolving around metabolic, mitochondrial and substrate handling terms (Fig. 4F-G), suggesting a coordinated dysregulation in skeletal muscle metabolism in c*Neb* KO mice. We also observed a significant increase in expression of myosin binding protein C1 (MYBPC1), myosin binding protein H (MYBPH) and ryanodine receptor 3 (RYR3) in the c*Neb* KO model (Supp. File 1). This is counterintuitive, given the contractile role of these proteins, but likely indicates a compensatory response in nemaline muscle to try and restore homeostasis.

**Figure 4:**
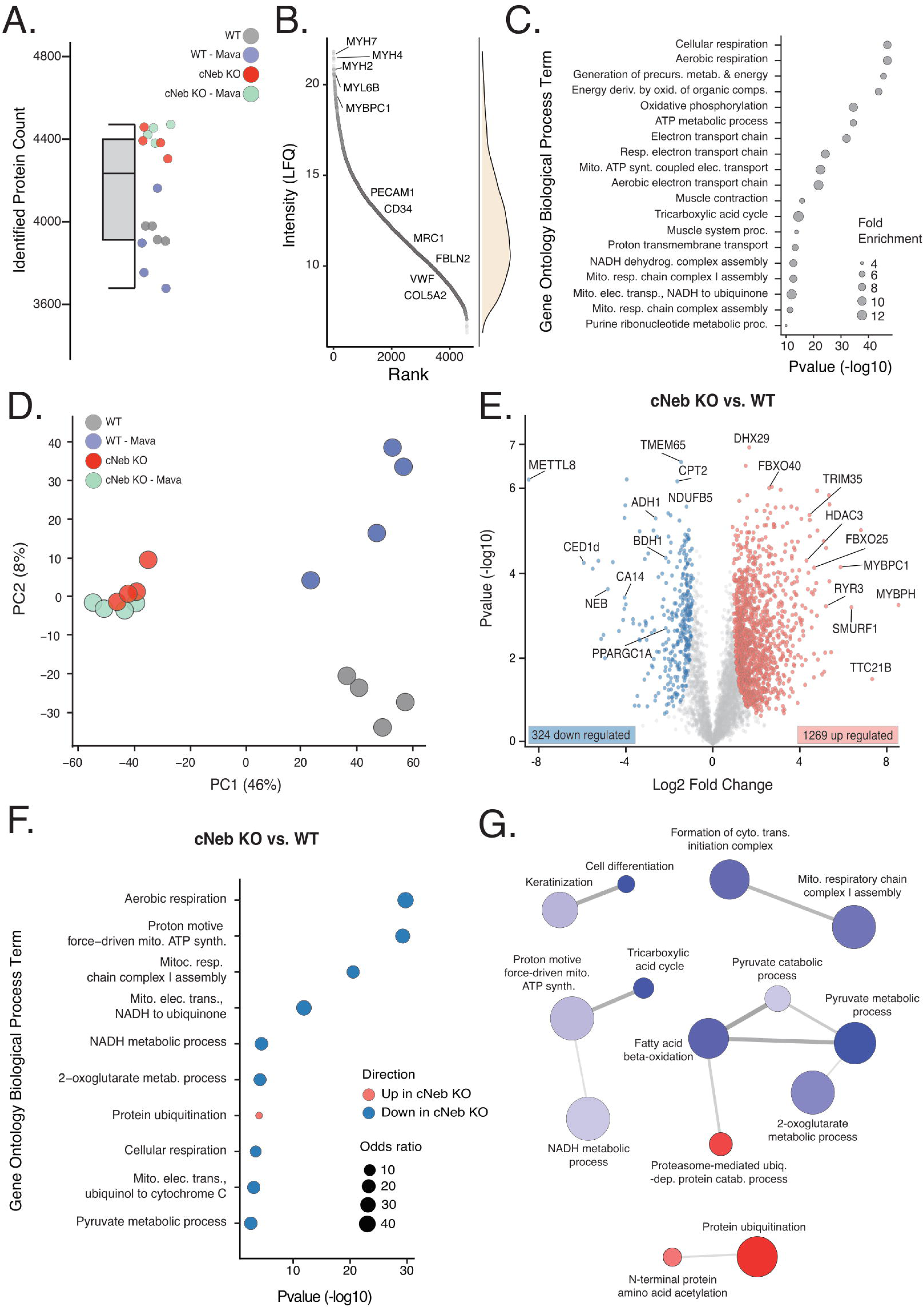
Global proteomic profiling of WT and cNeb KO mouse model. Number of identified protein hits (A) ranked by intensity (LFQ) with a density plot highlighting the distribution of proteins (B). Gene ontology biological processing terms associated from the top 250 ranked proteins highlighting the specificity of our global proteomic data from enriched myofibers (C). Principal component analysis (D) of all four conditions in our in-vivo mouse experiments, where each dot represents a single mouse. (E) volcano plot of protein abundances between cNeb KO and WT mouse soleus muscle highlighting significant differences between conditions (P < 0.05, FC > 1, see methods), where blue (down) and red (up) denote change in direction. These significant hits are used to identify enriched biological processing terms in (F), where the top 10 most enriched terms are represented (Supp File 3 for full list). Colour and size of each node denotes change in direction and odds ratio of term. (G). REVIGO (Supek *et al*., 2011) network of enriched biological processing terms of cNeb KO vs WT differential analyses. Nodes are connected if they annotate common protein, with the colour and weight of edges denoting strength of similarity. Network nodes colour represents directionality of comparison (cNeb KO vs WT, as per previous), strength in colour shows the number of annotations for GO term in underlying GOA data base (REVIGO), size is representative of P value (Supp File 3).

Our next comparison focussed on the impact of the four-week Mavacamten treatment in both WT and c*Neb* KO mice. Surprisingly, our initial PCA clustering analyses suggested that Mavacamten treatment had a relatively small effect on the soleus muscle of c*Neb* KO mice, but a larger impact on the proteome of WT muscle (Fig. 4D). Indeed, comparison of samples with and without Mavacamten treatment revealed 273 targets to be differentially regulated in WT myofibres (143 down-, 130 up-regulated; Fig. 5A; Supp. File 1), whereas 199 proteins were significantly differentially regulated following treatment in c*Neb* KO myofibres (97 down-, 102 up-regulated, Fig. 5B; Supp. File 1). Interestingly, very few proteins shared a common change in directionality across these comparisons, with only 5 proteins significantly increasing (including BRD3, HNRNPR and PKN2) and 6 decreasing in abundance (including BET1 and MRM1). Further analyses revealed no enrichment in biological processing meeting significance thresholds, for any of these identified protein lists. Taken together, interestingly, these data suggest that WT and c*Neb* KO mouse muscle fibres respond differently to short-term Mavacamten treatment.

**Figure 5:**
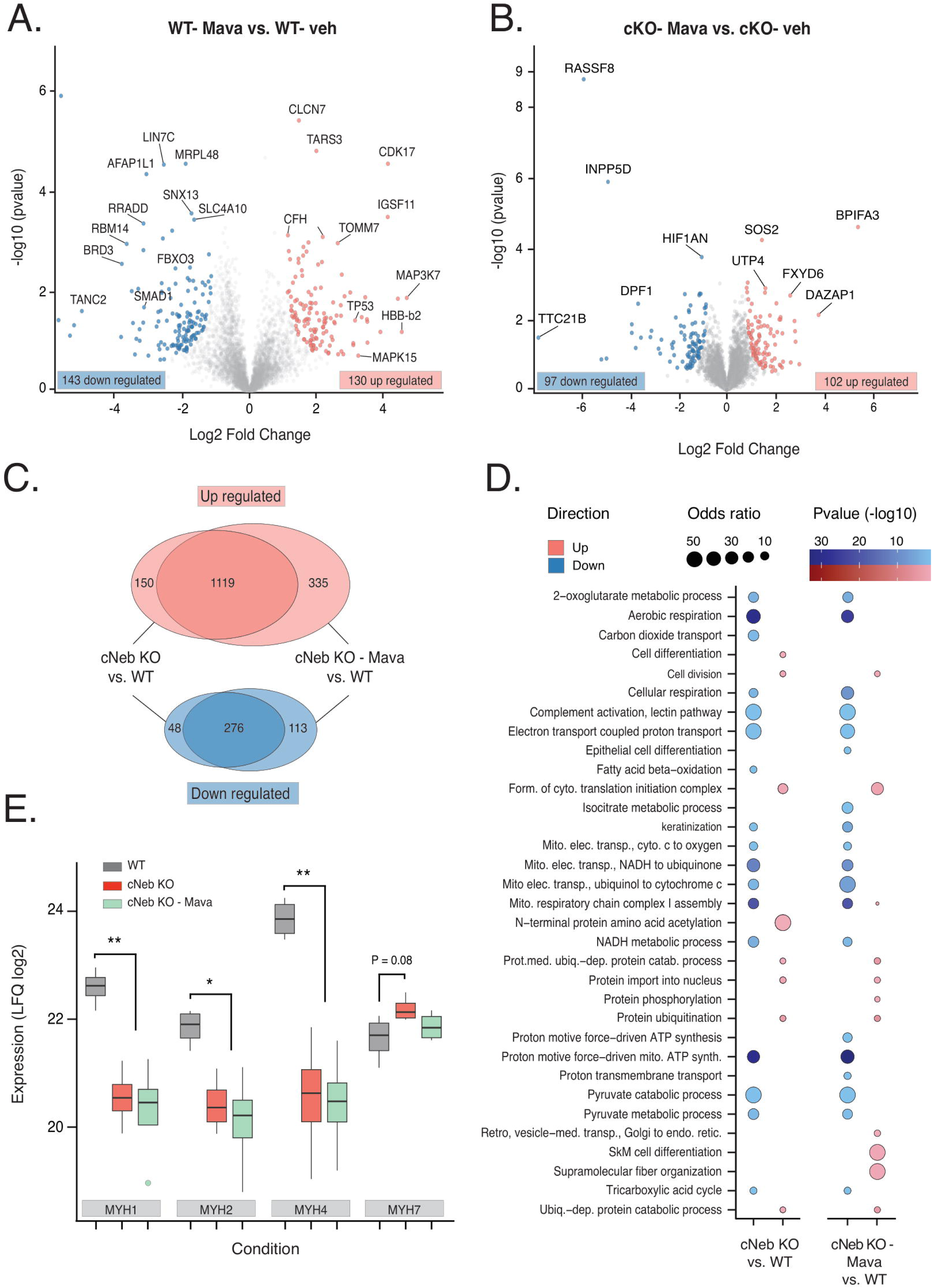
Proteomic abundance analysis investigating the impact of Mavacamten treatment on WT and cNeb KO mice. Volcano plot of protein abundances between ascertain the impact of 6-weeks of Mavacamten treatment in soleus muscle of WT (A) and cNeb KO (B) mice. Significant changes in protein abundance between conditions (P < 0.05, FC > 1, see methods) are highlighted by directionality of change (blue, decrease and red, increased in abundance). (C). Venn diagram displaying the number of significantly altered proteins between the comparisons of cNeb KO vs WT, and cNeb KO + Mavacamten vs WT. Increases in protein abundance within these comparisons are highlighted in red, decreases in blue (Supp Files 1 and 2). All significant protein hits between these comparison (e.g. cNeb KO and cNeb KO + Mavacamten, vs WT) used to compare enriched biological processing terms. Node colour shows change in direction of term in the specific comparison, strength of colour highlights Log10 P value and size of node represents the odds ratio of term. (E). Analysis of Log 2 transformed LFQ abundance of myosin isoforms (MYH1, 2, 4 and 7) as key determinants of fiber type in soleus mouse muscle. Box plots show median, inter-quartile range, range of points and outliers. N=4 for each condition, denoted by colour. Significant differences (ANOVA) at levels of P < 0.01 (*) and P < 0.001 (**).

To ascertain whether this Mavacamten treatment was able to restore any of the aberrant proteomic profile observed in the c*Neb* KO mouse model, we used the findings from our initial comparison (c*Neb* KO vehicle versus WT vehicle) as a baseline and compared these results with that of c*Neb* KO exposed to Mavacamten. Firstly, Mavacamten treatment in c*Neb* KO soleus myofibres (compared to WT vehicle) produced an enhanced number of proteins with modified expression (>1840 protein hits; FDR < 0.05), which is an increase of 185 upregulated proteins, and 65 down-regulated (Supp. File 1), compared with c*Neb* KO vehicle vs WT. Comparative analysis suggests that 150 proteins were significantly up-regulated in the c*Neb* KO vehicle muscle fibres, but not in the Mavacamten treated c*Neb* KO myofibres (both compared to WT vehicle as a control; Fig. 5C), leaving 335 novel protein hits that were upregulated in the Mavacamten-treated c*Neb* KO mouse muscle fibres (Fig. 5C). This same analysis in the opposite direction (down-regulated), showed 48 protein targets in our original comparison as significantly down-regulated (c*Neb* KO vehicle vs WT vehicle) but whose expression did not reach significance in any direction following Mavacamten treatment in c*Neb* muscle fibres (Fig. 5C). Of the 389 significantly down-regulated hits in the Mavacamten treated c*Neb* KO (WT vehicle as control), 113 were therefore newly identified. Gene ontology analysis of these protein hits reveals no coordinated or uniform biological processing term. No protein hits revealed in the control analysis (c*Neb* KO vehicle versus WT vehicle) reversed their directionality at a significant level following Mavacamten treatment. These data indicate that some protein targets, whose expression is dysregulated in c*Neb* KO mouse muscle fibres, may be recovered (but not reversed) to WT levels with Mavacamten treatment. We therefore sought to understand whether these targets were of relevance for the experimental model and treatment.

We re-analysed those gene ontology terms (biological process, Supp. File 3) that were most significantly enriched in the comparison of c*Neb* KO vehicle vs WT vehicle (Fig. 4F-G). Short-term treatment with Mavacamten was largely unable to recover or alter the dysregulation in these GO terms (Fig. 5D), with the most significantly enriched terms in our original comparison remaining significantly enriched in c*Neb* KO myofibres following Mavacamten treatment. Interestingly, c*Neb* KO mice exposed to Mavacamten did show an enrichment for the terms skeletal muscle cell differentiation and supramolecular fibre organization, possibly indicating a remodelling or recovery of fibre type distribution in the soleus muscle (Fig. 5D). However, analysis of myosin heavy chain protein isoform (MYH1, 2, 4 and 7) expression showed a significant reduction only in the expression of all fast type isoforms (MYH1, 2 and 4) compared to WT Vehicle control (Fig. 5E). Taken together, these data support that Mavacamten treatment is unable to adequately recover the metabolically dysregulated proteome of a c*Neb* KO mouse model of NM.

## Discussion

In the present study, we tested the potency of Mavacamten, a newly FDA-approved drug, *in vitro* and *in vivo*, using *NEB*-NM isolated single myofibres and a mouse model lacking nebulin (c*Neb* KO). Muscle fibres directly exposed to increasing concentrations of the compound showed promising results, with rescued proportions of myosin molecules in the ATP-conserving SRX state for both *NEB*-NM patients and c*Neb* KO mice. When the compound was given orally to the c*Neb* KO transgenic mice for a short period of time (four weeks), we observed a subtle proteome adaptation without any positive effects on metabolic proteins. Altogether, our findings highlight that Mavacamten is potent *in vitro,* whereas short-term *in vivo* administration may not be enough to rescue the metabolic *NEB*-NM phenotype.

### Potency of Mavacamten in vitro in NEB-NM

Here, promisingly, we show that the decreased level of myosin heads in the biochemical SRX state that occurs in *NEB*-NM patients as well as in c*Neb* KO mice (Ranu *et al*., 2022) can be fully rescued by Mavacamten at concentrations as low as 1 µM. This compound has already proven to be successful in treating dysfunctional myosin in hypertrophic cardiomyopathy (Toepfer *et al*., 2019). An analogue to this drug has also been beneficial for one class of skeletal myopathy due to *MYH7* mutations (Buvoli *et al*., 2024). Mavacamten is a small-molecule modulator of cardiac β/slow myosin heavy chain known to primarily reduce the steady-state ATPase activity by inhibiting the basal release rate of inorganic phosphate and ADP (Toepfer *et al*., 2019). More precisely, its presence in the active site of myosin heavy chain sub-fragment 1 promotes a closed state, stabilizing myosin heads in a folded-back conformation and subsequently the helical order filaments (Anderson *et al*., 2018). While our study using the Mant-ATP chase experiments focused on the biochemical myosin relaxed states, further structural work is needed to establish how Mavacamten and nebulin deficiency alter the structural Inter-Head-Motif-related folded-back conformation of myosin.

### Limited response of a four-week exposure to Mavacamten in cNeb KO mice

According to our principal component and subsequent proteomic analyses, soleus muscles from WT mice treated with Mavacamten at a dose of 2.5 mg.kg^-1^.day^-1^ for four weeks displayed clear effects. On the other hand, skeletal muscles from c*Neb* KO mice exposed to the same drug exhibited limited effects on the proteome. Interestingly, and in line with our finding, another recent study reveals that patients with obstructive hypertrophic cardiomyopathy have mixed Mavacamten responsiveness depending on their genotypes (Giudicessi *et al*., 2024).

Here, the differential response to Mavacamten between transgenic and control mice was further emphasized histologically (muscle fibre cross-sectional area). The limited effects on the proteome may be complex. It is tempting to speculate that such phenomenon is the consequence of a blunted activation of some of Mavacamten-related signalling pathways in the short term. These processes unlikely originate from a blockade of Mavacamten binding to myosin molecules as our *in vitro* data clearly demonstrate the potency of the drug in both WT and c*Neb* KO mice. To gain insights into other processes, we examined proteins that are up-regulated and down-regulated in WT animals with and without Mavacamten. Interestingly, when clustering the 273 targets that are differentially regulated (according to their biological families), many proteins related to ATP synthesis, glucose metabolism, lipid metabolism, mitochondrial morphogenesis, organisation and function are significantly different. As this metabolic reprogramming is not affected by the drug administration in c*Neb* KO mice, we speculate that the damages observed to metabolic proteins, organelles and/or related signalling pathways in this mouse model (Ranu *et al*., 2022; Tinklenberg *et al*., 2023) are not reversible, at least in the short term. Further studies are warranted.

## Conclusion

Mavacamten is clearly able to restore myosin ATP consumption *in vitro* in the context of NEB-NM. However, in contradiction with our initial hypothesis, a four-week administration of this drug in cNeb KO mice is not sufficient to rescue the muscle energy proteome. These results then highlight the need for future studies focusing on either drug delivery, higher dosage or longer treatment durations.

## Ethics approval and consent to participate

All human tissue was consented, stored, and used in accordance with the Human Tissue Act, UK, under local ethical approval (REC 13/NE/0373). All the mouse experiments were approved by the University of Arizona Institutional Animal Care and Use Committee (09-056) and were in accordance with the United States Public Health Service’s Policy on Humane Care and Use of Laboratory Animals.

## Availability of data and material

The mass spectrometry proteomics data have been deposited to the ProteomeXchange Consortium via the PRIDE partner repository (Perez-Riverol *et al*., 2022) with the dataset identifier PXD051963. All the data analyzed are also presented here in Fig. 1 to 4 and supplementary files.

## Authors’ contributions

JL, RAES, HG and JO contributed to the study conception and design. Material preparation, data collection and analysis were performed by JL, RAES, NR, JSK, TNB, CWP, NW, JV, JJV, EZ, JP, SH, HG and JO. The first draft of the manuscript was written by JL, RAES and JO and all authors commented on all versions of the manuscript and approved of the submitted version.

## Acknowledgements

We thank Thomas Nyegaard Beck for his assistance in many of the experiments outlined in the present manuscript. Additionally, we thank Dr Anders Karlsen for AI ilastik training and for developing the macro for CSA measurements with ImageJ (FiJi). Mass spectrometry analyses were performed by the Proteomics Research Infrastructure (PRI) at the University of Copenhagen (UCPH), supported by the Novo Nordisk Foundation (grant agreement number NNF19SA0059305).

